# Ketone Body Metabolism is Not Required for Improvement of Heart Failure by Ketogenic Diet in Mice

**DOI:** 10.1101/2024.08.30.610511

**Authors:** Zachary Foulks, Carla J. Weinheimer, Attila Kovacs, Jessica Michael, Kelly D. Pyles, Thiago N. Menezes, Kevin Cho, Gary J. Patti, Kyle S. McCommis

## Abstract

Failing hearts increasingly metabolize ketone bodies, and enhancing ketosis improves heart failure (HF) remodeling. Circulating ketones are elevated by fasting/starvation, which is mimicked with a high-fat, low-carbohydrate “ketogenic diet” (KD). While speculated that KD improves HF through increased ketone oxidation, some evidence suggests KD paradoxically downregulates cardiac ketone oxidation despite increased ketone delivery. We sought to clarify the significance of cardiac ketone metabolism during KD in HF. Mice were subjected to transverse aortic constriction with apical myocardial infarction (TAC-MI) and fed either low-fat (LF) control or KD. Cardiac-specific mitochondrial pyruvate carrier 2 (csMPC2-/-) mice were used as a second model of heart failure. In both mice, feeding a KD improved HF, determined by echocardiography, heart weights, and gene expression analyses. Although KD increases plasma ketone bodies, gene expression for ketone metabolic genes is decreased in the hearts of KD-fed mice. Cardiac-specific β-hydroxybutyrate dehydrogenase 1 (csBDH1-/-), the first enzyme in ketone catabolism, mice were also studied and crossed with the csMPC2-/-mice to create double knockout (DKO) mice. These mice were aged to 16 weeks and switched to LF or KD, and KD was able to completely normalize the hearts of both csMPC2-/- and DKO mice, suggesting that ketone metabolism is unnecessary for improving heart failure with ketogenic diet. These studies were then repeated, and mice injected with U-^13^C-β-hydroxybutyrate to evaluate ketone metabolism. KD feeding significantly decreased the enrichment of the TCA cycle from ketone body carbons, as did the BDH1-deletion in DKO mice. Gene expression and respirometry suggests that KD instead increases cardiac fat oxidation. In conclusion, these results suggest that ketogenic diet decreases cardiac ketone metabolism and does not require ketone metabolism to improve heart failure.

## introduction

Compared to healthy hearts, failing hearts exhibit decreased fat oxidation and enhanced glycolysis for ATP synthesis. Failing hearts also increasingly metabolize ketone bodies,^1^ and enhancing ketosis improves heart failure (HF) remodeling.^1-3^ Circulating ketones are elevated by fasting/starvation, which is mimicked with a high-fat, low-carbohydrate “ketogenic diet” (KD). Previous studies described cardioprotective effects of KD,^1, 2^ and ongoing clinical trials are investigating its cardiovascular benefits. While speculated that KD improves HF through increased ketone oxidation, some evidence suggests KD paradoxically downregulates cardiac ketone oxidation despite increased ketone delivery.^2, 4, 5^ We sought to clarify the significance of cardiac ketone metabolism during KD in HF.

WT mice were subjected to transverse aortic constriction with apical myocardial infarction (TAC-MI) to induce HF.^1^ After two weeks of recovery and cardiac remodeling, echocardiography was performed and mice switched to low-fat diet (LF; 70/10/20% kcal carbohydrate/fat/protein; Bio-Serv F1515) or KD (1.8/93.4/4.7% kcal carbohydrate/fat/protein; Bio-Serv F3666). After two weeks, echocardiography was repeated and hearts collected. KD prevented further increase in end-systolic (ESV) and end-diastolic volumes (EDV) and maintained ejection fraction (EF) compared to LF (Figure [A]), suggesting improved dilated cardiomyopathy. Indeed, KD-feeding significantly decreased expression of heart failure and fibrosis genes (Figure [B]). Despite KD-feeding increasing ketosis, the hearts decreased expression of ketolytic enzymes β-hydroxybutyrate dehydrogenase 1 (*Bdh1*) and 3-oxoacid CoA-transferase 1 (*Oxct1*), and upregulated expression of fat oxidation genes (Figure [B]).

**Figure.**
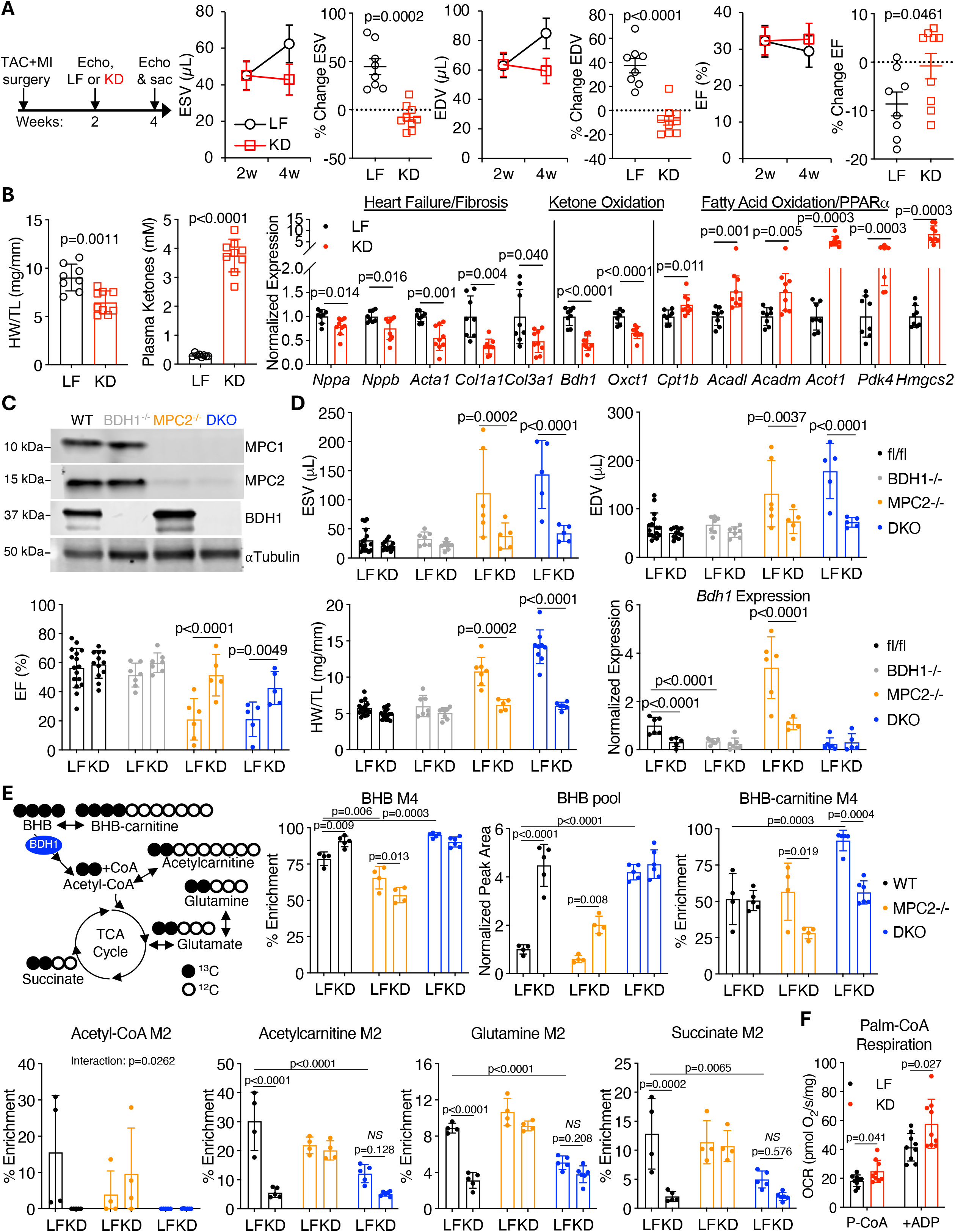
Ketone body metabolism is not required for ketogenic diet improvement of heart failure. **A**, Echocardiography measures of end-systolic volume (ESV), end-diastolic volume (EDV), and ejection fraction (EF) of female C57BL/6J mice subjected to transverse aortic constriction with apical myocardial infarction (TAC-MI) 2 weeks post-surgery, and after 2 weeks of low-fat (LF) or ketogenic diet (KD) feeding (4-weeks post-surgery). Data also expressed as %-change from 2-to 4-weeks (n=8-9 per group). **B, Left**, Heart weight normalized to tibia length (HW/TL) and total plasma ketone concentrations of mice subjected to TAC-MI and fed LF or KD (n=8-9 per group). **B, Right**, Cardiac gene expression for heart failure and fibrosis genes, ketone metabolism genes, and fat oxidation genes (n=8-9 per group). **C**, Representative western blot showing cardiac deletion of BDH1 and/or MPC1/2 proteins from csBDH1-/-, csMPC2-/-, and DKO hearts. **D**, ESV, EDV, EF, HW/TL, and *Bdh1* expression from WT (*Mpc2* and *Bdh1* floxed), csBDH1-/-, csMPC2-/-, and DKO mice fed LF or KD (n=5-17 per group). **E**, Schematic of U-^13^C-β-hydroxybutyrate (BHB) labeling, and the primary enrichment products for BHB, BHB-carnitine, acetyl-CoA, acetylcarnitine, glutamine, and succinate, as well as BHB pool size (n=4-6 per group). **F**, Oxygen consumption rates (OCR) of cardiac muscle fibers from LF of KD-fed mice when provided palmitoyl-CoA + carnitine, and then adenosine diphosphate (ADP) (n=9 per group). Data expressed as mean ± SD and analyzed by Student’s 2-tailed *t* test (**A, B**, and **F**), or 2-way ANOVA with Tukey multiple comparison’s test (**D** and **E**) with GraphPad Prism v10.3.0.

Similar improvements in remodeling from TAC-MI were observed previously;^1^ however, KD was initiated prior to surgery in that study. These results suggest KD improved HF despite potentially decreased cardiac ketone oxidation.

An appreciable role for ketone oxidation benefitting HF still could not be discounted.

Therefore, we employed cardiac-specific BDH1 knockout mice (csBDH1-/-),^1^ deleting the enzyme converting β-hydroxybutyrate (BHB), the primary circulating ketone, to acetoacetate for further oxidation. We also crossed these mice with cardiac-specific mitochondrial pyruvate carrier 2 (csMPC2-/-) mice,^2^ which spontaneously develop HF, to create MPC2/BDH1 double knockout mice (DKO) (Figure [C]). Mice consumed normal chow until 16 weeks of age, then LF or KD for 3 weeks. csBDH1-/-mice displayed normal ESV, EDV, and EF on either diet (Figure [D]). csMPC2-/-hearts were dilated with reduced EF on LF, but were normalized by KD.^2^ DKO hearts demonstrated similar remodeling on LF, with KD normalizing their dilation, contraction, and weight. HF and fibrosis gene expression was normalized by KD in both csMPC2-/- and DKO hearts (data not shown). Lastly, while the failing hearts of LF-fed csMPC2-/- mice upregulated *Bdh1*, KD-feeding reduced *Bdh1* expression in these and WT hearts (Figure [D]). csBDH1-/- and DKO hearts expressed little *Bdh1*, regardless of diet. These results suggest KD reduces cardiac ketone metabolism, and rescuing HF with KD does not require BDH1.

To evaluate the fate of cardiac BHB, we placed 16-week-old WT, csMPC2-/-, and DKO mice on LF or KD for 2 weeks. We then i.p. injected 1 mg/g U-^13^C-BHB, and anesthetized the mice with 1-2% inhaled isoflurane and freeze clamped the beating hearts *in situ* 30 min post injection. Metabolites were extracted from the hearts and UHPLC-MS performed. In WT hearts, KD increased uptake of fully-labeled BHB and increased the pool size of BHB without altering enrichment of BHB-carnitine, which is formed separately from the ketone oxidation pathway (Figure [E]). Enrichment of ^13^C-BHB carbons into acetyl-CoA, acetylcarnitine, glutamine, and succinate were all reduced on KD in WT hearts (Figure [E]), indicating reduced ketone oxidation. In csMPC2-/-hearts, BHB pool size and enrichment were decreased versus WT.

However, in these hearts KD did not alter enrichment of BHB oxidation products, likely due to KD decreasing *Bdh1* expression to normal levels in csMPC2-/-hearts (Figure [D]).^2^ In DKO hearts, BHB pool size/enrichment and BHB-carnitine enrichment were elevated, suggesting reduced ketone oxidation. Enrichment of acetyl-CoA and TCA cycle metabolites also decreased in DKO hearts (Figure [E]), and KD had no significant effect besides normalizing enrichment of BHB-carnitine. These results confirm that cardiac ketone metabolism decreases with KD-feeding or BDH1 deletion.

Since KD reduces ketone metabolism and increases expression of fat oxidation genes (Figure [B]),^4^ we next tested whether KD enhanced fat oxidation. WT mice were fed LF or KD for 2 weeks, then mitochondrial respiration was assessed in cardiac muscle fibers with palmitoyl-CoA + carnitine.^2^ Palmitoyl-CoA-supported respiration was increased in fibers from KD versus LF-fed mice (Figure [F]). Thus, KD decreases cardiac ketone metabolism and enhances fat oxidation.

In conclusion, KD consumption decreases cardiac ketone oxidation due to reduced *Bdh1* and *Oxct1* expression.^4,5^ As hearts deficient in *Bdh1* are rescued by KD, this suggests ketone metabolism is not required for this effect. Instead, KD likely improves HF by enhancing fat oxidation. As cardiac BHB delivery increases on KD, it is also possible that non-metabolic effects of BHB, such as modifying proteins by lysine β-hydroxybutyrylation, could play a role. Ongoing studies will assess the importance of these modifications in HF.

## Acknowledgements

We thank Dr. Daniel P. Kelly from the University of Pennsylvania for providing the BDH1 floxed mice.

